# A Neural Oscillatory Signature of Reference

**DOI:** 10.1101/072322

**Authors:** Mante S. Nieuwland, Andrea E. Martin

## Abstract

The ability to use words to refer to the world is vital to the communicative power of human language. In particular, the anaphoric use of words to refer to previously mentioned concepts (antecedents) allows dialogue to be coherent and meaningful. Psycholinguistic theory posits that anaphor comprehension involves reactivating a memory representation of the antecedent. Whereas this implies the involvement of recognition memory, the neural processes for reference resolution are largely unknown. Here, we report time-frequency analysis of four EEG experiments to reveal the increased coupling of functional neural systems associated with referentially coherent expressions compared to referentially ambiguous expressions. Despite varying in modality, language, and type of referential expression, all experiments showed larger gamma-band power for coherence compared to ambiguity. Beamformer analysis in high-density Experiment 4 localised this increase to posterior parietal cortex around 400-600 ms after anaphor-onset and to frontal-temporal cortex around 500-1000 ms. We argue that the observed gamma-band power increases reflect successful referential binding and resolution, which links incoming information to antecedents through an interaction between the brain’s recognition memory networks and frontal-temporal language network. We integrate these findings with previous results from patient and neuroimaging studies, and outline a nascent cortico-hippocampal theory of reference.

---

Reference, or the ability to link linguistic representations to the world, makes language an immensely powerful tool for communication. Reference gives language intentionality, or ‘aboutness’, whether reference is made to the real world or to a hypothetical one. This essential computation presents a complex problem that dominates the philosophy of language to this day (e.g., Recanati 1993). Cognitive science shows that the crucial role of reference starts as early as language development itself, when children learn the meaning of words through understanding the referential intention of a speaker (e.g., Bloom 2000). From this perspective, *words* are the spoken or written symbols that people use to denote referents in the physical world around them, or in the conversations and stories that they engage in. Grasping a word’s referential meaning is therefore a key challenge when establishing the intended meaning of a speaker. Moreover, in connected text or dialogue, the referential meaning of a word frequently often involves reference to a previously mentioned concept, such that the comprehender must establish a referential relationship between a given referring expression and its antecedent. Thus, the computational problem that reference presents can be, at least in some sense, boiled down to relating two representations that are separated from each other in time and/or by other representations.

Given these circumstances, how is reference computed in the mind and brain? Psycholinguistic theories of reference have a broad consensus on the involvement of memory in referential processing, which we will discuss shortly, but a neurobiological mechanism for reference is still largely unknown. In fact, as of 2016, neurobiological accounts of sentence-level language comprehension have not yet been articulated to the level of reference (e.g., Friederici 2012; Price 2012; Bornkessel-Schlesewky and Schlesewsky 2013; Hagoort and Indefrey 2015), probably because these models primarily focus on accounting for syntactic and semantic processing phenomena.

To acquire data relevant to this theoretical gap, the current study performs oscillatory analysis of four encephalography (EEG) experiments on the neural signature of reference (Experiment 1: Nieuwland and Van Berkum 2006; Experiment 2: Nieuwland, Otten and Van Berkum 2007; Experiment 3: Martin, Nieuwland and Carreiras 2012; Experiment 4: Nieuwland 2014). Crucially, these experiments vary in language (Dutch, Spanish or English), modality (written sentences or spoken story comprehension) and manipulated linguistic expression (pronouns, noun phrases, ellipsis). However, their theoretical aim was similar: all the experiments examined how people understand expressions that refer back to a previously mentioned concept, and, more specifically, compared expressions with a straightforward, coherent referential meaning to expressions that are referentially ambiguous or otherwise referentially insufficient. Through oscillatory analysis on these datasets, we revealed the increased coupling of functional neural systems associated with referential coherence compared to referential ambiguity. Moreover, Experiment 4 involved high density EEG recording, which allowed us to localize the source of this increased neural coupling using a Beamformer procedure (Gross et al. 2001).

## A computational architecture for reference

A word that refers back to a previously mentioned concept (the antecedent) is an *anaphor*. The “memory-based” processing literature on text anaphora argues that anaphor comprehension involves the reactivation of the antecedent from a memory representation of the discourse (e.g., Dell, McKoon and Ratcliff 1983; Gernsbacher 1989; Gerrig and McKoon 1998; see also Sanford and Garrod 1989, 2005), followed by the subsequent integration of the antecedent into the overall representation of the narrated event. Psycholinguistic experiments have demonstrated that this process proceeds very rapidly when the anaphor shares sufficient semantic and syntactic features with the antecedent (for a review, see Garnham 2001; Sturt 2013). Feature-based antecedent reactivation enables the recognition of the antecedent, and therefore the establishment of a referential link between multiple instantiations of the same concept despite linguistic form differences, such that anaphora are not understood as mere repetitions of the antecedent (e.g., “The old man laughed. The man/Peter/he was happy”, see Garrod, Freudenthal and Boyle 1994; Almor 1999).

In our view, the abovementioned broad strokes processing framework on antecedent reactivation equates to, or, minimally, involves recognition memory function. Such a hypothesis is consistent with a broader approach to linguistic dependency resolution known as the cue-based retrieval framework (CBR; Lewis et al. 2006; McElree 2006; Martin and McElree 2008, 2009; Martin 2016). CBR builds on the computational architecture of human recognition memory and extends its mechanistic principles to language processing contexts. In the CBR framework, memory representations like antecedents are organised and recovered by virtue of their content (*content-addressable*), and they are elicited directly, without a so-called “search” through memory, based on the contact of its features with memory cues that are available in the anaphor (*direct-access, cue-based retrieval*). The extent to which unintended memory representations with content-overlap interfere with this process (*cue-based retrieval interference*) naturally emerges as the primary determinant of retrieval difficulty (e.g., Nairne 2002; Öztekin and McElree 2007).

This cognitive-mechanistic account places anaphoric reference as a systemic computation in a larger processing model of language (e.g., Martin 2016) and can lay the groundwork for developing an account of its neurobiology. This account, as do related memory-based accounts (Sanford and Garrod 1989; Gerrig and McKoon 1998), also offers a straightforward explanation for observed differences in the ease of anaphor comprehension, including differences between referentially coherent and ambiguous expressions as used in the current study. Relative to referentially coherent expressions, ambiguous expressions are associated with greater retrieval interference, which depends on the degree of content-overlap between anaphor and the intended antecedent relative to the content-overlap between the anaphor and other memory representations (Martin and McElree 2008, 2009, 2011; Van Dyke and McElree 2011). This explains why ambiguous expressions, whose features fail to elicit a unique target from memory, result in slower processing times and delayed comprehension (e.g., Gernsbacher 1989; MacDonald and MacWhinney 1990; Stewart, Holler & Kidd, 1990).

## Neurobiological implications of memory-based anaphor resolution

Memory-based theories of anaphor resolution, including the CBR account, also harbour predictions, albeit only implicitly, regarding underlying neural processes. In recognition memory research, successful retrieval (i.e., recognition) is associated with increased activity in and connectivity between the medial temporal lobe and posterior parietal cortex (Shannon and Buckner 2004; Gonzalez et al. 2015; for a review, see Wagner, Shannon, Kahn, and Buckner 2005). This may reflect the reactivation of, and bringing back into the focus of attention information that was previously encoded by the hippocampal system (e.g., McClelland, McNaughton and O’Reilly 1995; Öztekin, McElree, Staresina and Davachi 2009; Staresina, Henson, Kriegeskorte and Alink 2012; Levy and Wagner 2013; Gordon, Rissman, Kiani and Wagner 2013). In light of this literature, we predict that similar patterns of brain activity may occur if anaphora are understood through the retrieval or reactivation of an antecedent through recognition memory.

Consistent with this first prediction, Nieuwland, Petersson and Van Berkum (2007) observed BOLD activity increases in the hippocampus for pronouns that matched a unique antecedent in the sentence relative to pronoun that did not. Furthermore, damage to the hippocampus, which leads to hippocampal amnesic syndrome, is associated with impairments in pronoun production and comprehension (e.g., MacKay, James, Taylor and Marian 2007; Kurczek, Brown-Schmidt and Duff 2013; see also Almor, Kempler, MacDonald, Anderson and Tyler 1999, for related findings on pronoun comprehension in Alzheimer’s disease), and with reduced used of definite reference to previously discussed items (Duff, Gupta, Hengst, Tranel and Cohen 2011).

A second prediction involves the brain’s neural oscillatory activity, or the synchronization of its neural populations as measured in electrical and magnetic activity (EEG, MEG, ECoG). Neural oscillations reflect the transient coupling or uncoupling of functional neural systems, or cell assemblies (e.g., Engel, Fries and Singer 2001; Buzsaki and Draghun 2004), thereby offering a window into the functional network dynamics of human cognition. Oscillatory activity in particular the theta-frequency (3-8 Hz) and gamma-frequency band (> 30 Hz) increases in power for successful recognition (e.g., Herrmann, Munk and Engel 2004; Mormann et al. 2005; Jacobs, Hwang, Curren and Kahana 2006; Osipova, Takashima, Oostenveld, Fernandez, Maris and Jensen 2006; Jensen, Kaiser, and Lachaux 2007; Burke et al. 2014; Gonzalez et al. 2015; for a review, see Bastiaansen and Hagoort 2003; Düzel, Penny and Burgess 2010; Klimesch, Freunberger and Sauseng 2010; Nyhus and Curran 2011; Lisman and Jensen 2013). Like successful memory recognition, successful anaphor resolution may lead to an increase in gamma-band and/or theta activity. Consistent with this prediction, unpublished data from Van Berkum, Zwitserlood, Bastiaansen, Brown and Hagoort (2004) showed a gamma-band increase around 40 Hz for referentially coherent pronouns compared to mismatching pronouns. The current study follows up on this work and tests the prediction that referentially coherent expressions elicit increased gamma/theta-band oscillatory activity compared to referentially ambiguous expressions.

The observation that referentially coherent expressions elicit more gamma/theta oscillatory activity than referentially problematic expressions, and not the other way around, would deviate in a crucial way from the commonly observed pattern that problematic utterances lead to increased brain activity compared to unproblematic utterances. Research on the neurobiology of language comprehension often relies on violation-paradigms (e.g., syntactically or semantically anomalous sentences), yielding conclusions about language comprehension based on the increase in activity when language processing is atypical or fails (e.g., Nieuwland, Petersson and Van Berkum 2007; Nieuwland, Martin and Carreiras 2012). This approach has been generally very fruitful, but its conclusions about ‘normal’ language comprehension are inherently limited. Therefore, finding increased oscillatory brain activity for successful reference, across different modalities and linguistic manipulations, offers an important initial step in describing the common processes involved in reference.

Such a finding on its own does not constitute unequivocal, direct support for the involvement of recognition memory networks during anaphora resolution. Changes in gamma/theta oscillatory activity are not unique to recognition memory, and have been observed for different language comprehension processes (e.g., Roehm, Schlesewsky, Bornkessel, Frisch and Haider, 2004; Davidson and Indefrey 2007; Wang, Zhu and Bastiaansen 2012; Rommers, Dijkstra and Bastiaansen 2013; Bastiaansen and Hagoort 2015; Lewis, Wang and Bastiaansen 2015; Lam, Schoffelen, Udden, Hulten and Hagoort 2016). Lewis and Bastiaansen (2015) proposed that gamma-band activity during sentence comprehension indexes predictive processing, and that theta-band activity indexes lexical-semantic retrieval. Such a frequency-to-function mapping, while intriguing, risks oversimplification of dynamic brain activity, and we will return to this issue in our discussion.

To obtain additional support for our hypothesis about the involvement of recognition memory during anaphor resolution, we therefore investigated whether oscillatory activity increases are at least in part generated by brain regions strongly associated with recognition, like the medial temporal lobe and posterior parietal cortex. Importantly, anaphor comprehension requires antecedent reactivation but also the integration of this information into the unfolding sentence context (e.g., Sanford and Garrod 1989), and therefore also relies ongoing language processes (e.g., syntactic structure building, lexical-semantic processing). Therefore, we expected to see additional involvement of the traditional frontal-temporal language network (e.g., Hagoort 2013; Hagoort and Indefrey 2014; Fedorenko and Thompson-Schill 2015; Friederici and Singer 2015).

## The current study

The current study seeks to observe a signature of reference resolution in brain oscillations. We compared oscillatory activity induced by referentially coherent expressions with activity induced by referentially ambiguous expressions, using existing data from four previously reported event-related brain potential (ERP) experiments. These experiments differed in the language of study, modality and the referential expression of interest.

Experiment 1 (Nieuwland and Berkum 2006) examined the comprehension of written Dutch sentences with pronouns that matched either one or two characters (e.g., “John told Mary/David that he needed to buy insurance”). Experiment 2 (Nieuwland et al. 2007) examined the comprehension of spoken Dutch stories with noun phrase anaphora that matched either one or two characters (e.g., ‘the nephew’ in a context with one or two nephews). Experiment 3 (Martin et al. 2012) examined the comprehension of written Spanish sentences with noun phrase ellipsis that matched or mismatched its antecedent in gender (e.g., “la camiseta … otra/otro”, with otra/otro being the feminine or masculine gender equivalent of ‘another’; see also Martin, Nieuwland & Carreiras, 2014). Experiment 4 (Nieuwland 2014) examined the comprehension of written English sentences with pronouns that matched or mismatched the only mentioned character in the sentence (e.g., “John said that he/she was a very happy person”).

Despite the differences between the experiments, referentially ambiguous expressions in each experiment elicit a slowly unfolding, sustained frontal negativity in the ERP waveform^1^ compared to referentially coherent expressions (see Nieuwland and Van Berkum 2008, for a review), suggesting the involvement of qualitatively similar processes. Here, we predicted that oscillatory activity in these experiments would show a gamma/theta-band increase for coherence compared to ambiguous expressions, and that at least some of the observed differences would generate from the recognition memory network (posterior parietal cortex) and the frontal-temporal language network.

## METHODS AND MATERIALS

Table 1 shows example materials from each experiment and a brief description of the relevant linguistic manipulation. Note that in the four experiments reported here, participants also read or listened to a large number of filler sentences along with the experimental sentences described here. Full methodological details regarding participants, materials and procedure are available in the previously published report for each experiment. Here we describe the methods that are relevant to the current analyses.

**Table 1.**
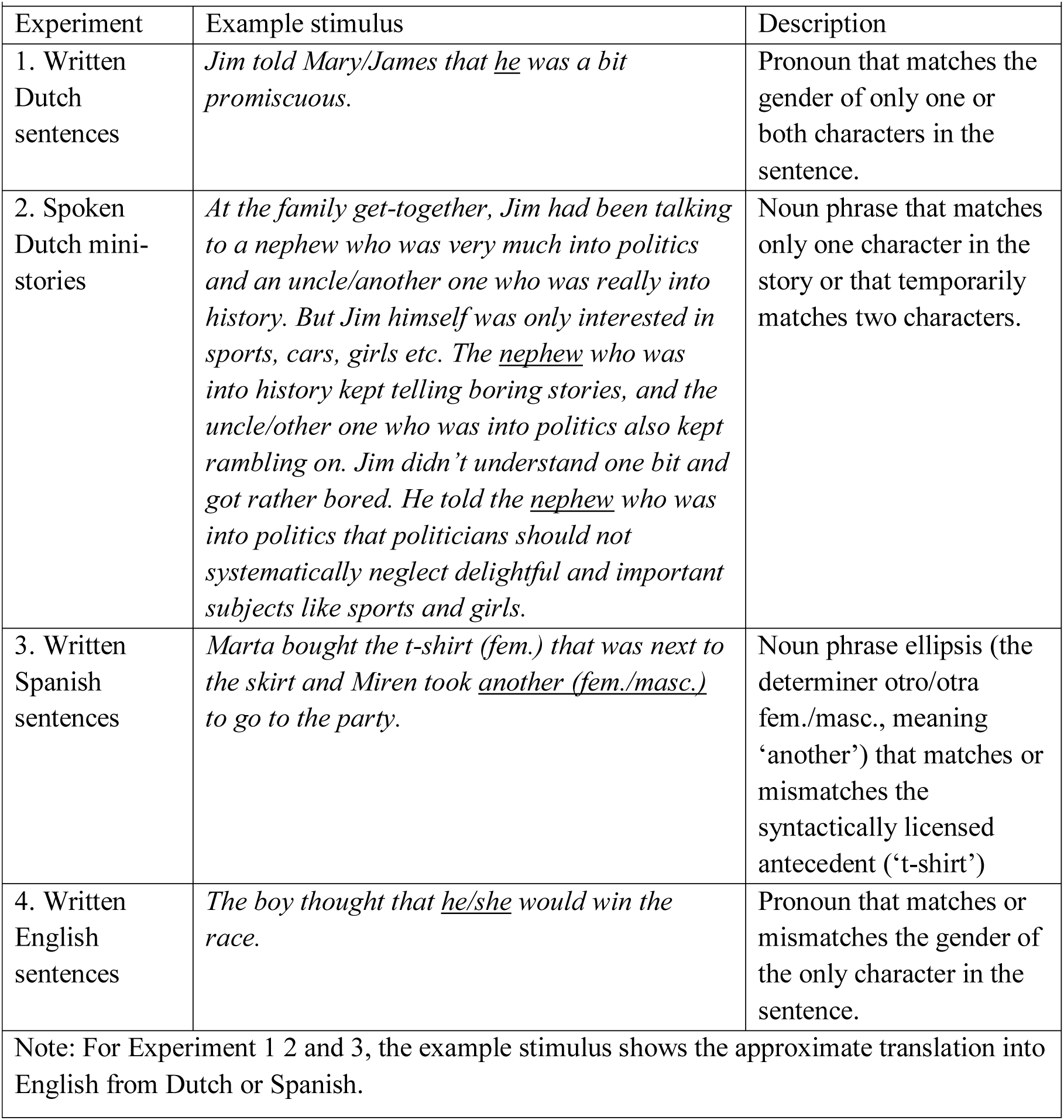
Example materials for each of the four experiments. Critical words are underlined for expository purposes only. Each example shows the referentially coherent/ambiguous version separated by a slash character.

### Participants, Materials and Procedure

In Experiment 1 (Nieuwland and Van Berkum 2006), thirty-two native speakers of Dutch read 120 grammatically correct Dutch sentences that described an interaction between two individuals. Sixty sentences described two individuals of different gender and contained a referentially coherent pronoun that matched exactly one individual (e.g., “Anton forgave Linda because she …”), and the other 60 sentences described two individuals of the same gender and contained a referentially ambiguous pronoun that matched both individuals (e.g., “Mary forgave Linda because she …”). Participants read the sentences one word at a time from the centre of a display, with word duration dependent on word length. Participants did not perform a secondary task.

In Experiment 2 (Nieuwland et al. 2007), thirty-one native speakers of Dutch listened to 90 naturally spoken Dutch mini-stories of 5 sentences that described a scenario with one protagonist and two secondary characters. The two secondary characters were always denoted with a noun phrase followed by a relative clause of at least four words, and they could be denoted with different noun phrases or with the same noun phrase. When the same noun phrase was used, the last word of the relative clause disambiguated the temporarily ambiguous expression (e.g., “the nephew who was really into politics/sports”). The third and the fifth sentence of each story mentioned one of the secondary characters using a referential expression that was either referentially coherent or that was temporarily ambiguous. In the current analysis, we excluded stories in which the third sentence contained an ambiguity but in which the fifth sentence was referentially coherent, in order to match ambiguous and coherent conditions on story position. This selection resulted in a maximum of 60 trials per condition. Participants did not perform a secondary task.

In Experiment 3 (Martin et al. 2012), twenty-two native speakers of Spanish each read 60 Spanish sentences in which the gender of the critical word (otro or otra; the remaining determiner of the elided noun phrase) was grammatically correct or incorrect given that of the antecedent (e.g., ‘camiseta’ in “Marta se compró la camiseta que estaba al lado de la falda y Miren cogió otra/otro para salir de fiesta”, see Table 1 for approximate translation into English). The critical word was always followed by at least three other words. Participants read the sentences one word at a time from the centre of the display and answered intermittent comprehension questions about the presented sentences throughout the experiment.

In Experiment 4 (Nieuwland 2014), nineteen native speakers of English each read 180 grammatically correct English sentences that introduced a female or male character, followed by ‘verb-ed that’ and subsequently a male of female pronoun and another 4 words. Ninety sentences contained a referentially coherent pronoun that matched the character and 90 sentences contained a referentially ambiguous pronoun that did not match the character (e.g., “Clifford mentioned that he/she was getting a divorce.”). Participants read the sentences one word at a time at a pace of 2 words per second, and were not asked to perform any secondary task.

### EEG data recording and pre-processing

In Experiment 1, continuous EEG data was collected from 30 standard channels (10/20 system) using an ActiCap (Brain Products, Munich, Germany), plus two additional EOG electrodes. The EEG was recorded with a 5-s time constant and a 100-Hz low pass filter, and sampled at 500□Hz. All electrode impedances were kept below 5□kΩ. In this and all following experiments, we used Brain Vision Analyzer software 2.0 (Brain Products) to pre-process the raw EEG data. The EEG data were re-referenced off-line to the average of both mastoids, and filtered with a 1-Hz high pass filter (48 dB/Oct). Data segments from −1 to 2.5□s relative to the onset of the pronoun in each sentence were extracted and corrected for ocular artefacts using a method based on Independent Component Analysis. After that, we applied automatic artifact rejection based on three rejection criteria simultaneously: an amplitude criterion of ±90□μV, a gradient criterion (i.e., the maximum admissible voltage step between two adjacent time points) of 50□μV, and a difference criterion (i.e., the maximum admissible absolute difference between two values within each EEG epoch) of 120□μV. Only participants with at least 40 trials in each of the conditions were included, leaving 28 participants for analysis. The referentially coherent and ambiguous conditions retained on average 54 and 55 trials per subject, respectively.

In Experiment 2, EEG data recording and pre-processing was identical to that in Experiment 1. Only participants with at least 40 trials in each of the conditions were included, leaving 26 participants for analysis. The referentially coherent and ambiguous conditions retained on average 53 and 52 trials per subject, respectively.

In Experiment 3, EEG data was collected from 29 standard channels (10/20 system) using an ActiCap (Brain Products, Munich, Germany), plus four additional EOG electrodes. The EEG was recorded continuously with a 10-s time constant and a 100-Hz low pass filter, and sampled at 250□Hz. The pre-processing procedure was identical to that of Experiment 1 and 2. Only participants with at least 20 trials in each of the conditions were included, leaving 18 participants for analysis. The two conditions had the same average number of trials (28) per subject entering the analysis.

In Experiment 4, EEG data was recorded at a sample rate of 512 Hz and with 24-bit AD conversion using the Biosemi ActiveTwo system (BioSemi BV, Amsterdam, The Netherlands). This system’s hardware is completely DC coupled and applies digital low pass filtering through its ADC’s decimation filter (the hardware bandwidth limit), which has a 5^th^-order sinc response with a -3 dB point at 1/5th of the sample rate (i.e., approximating a low-pass filter at 100□Hz). Data was recorded from 64 EEG, 4 EOG, and 2 mastoid electrodes using the standard 10/20 system. The pre-processing procedure was identical to that of Experiment 1-3 except for the automated artifact rejection procedure. A slightly more liberal artefact criterion was used to retain enough participants for analysis, using an amplitude criterion of ±100 μV, a gradient criterion (i.e., the maximum admissible voltage step between two adjacent time points) of 50 μV, and a difference criterion (i.e., the maximum admissible absolute difference between two values within each EEG epoch) of 150□μV. Only participants with at least 50 trials in each of the conditions were included, leaving 16 participants for analysis. The two conditions had the same average number of trials (70) per subject entering the analysis.

### Time-frequency analysis

Time-frequency analysis was performed following the same procedure in Experiment 1 to 4, using the Fieldtrip software package (Oostenveld, Fries, Maris and Schoffelen 2011). To optimize the trade-off between time and frequency resolution, time-frequency analysis was performed using the extracted epochs in two different, partially overlapping frequency ranges. In the low frequency range (2–30□Hz), a 400-ms Hanning window was used to compute power changes in frequency steps of 1□Hz and time steps of 10□ms. In the high frequency range (25–90□Hz), time-frequency analysis was performed using a multitaper approach (Mitra and Pesaran 1999), computing power changes with a 400-ms time-smoothing and a 5-Hz frequency-smoothing window, in 2.5□Hz frequency steps and 10□ms time steps. Subsequently, power changes per trial in the post-stimulus interval were computed as a relative change from a baseline interval spanning from −0.5 to −0.3□s relative to critical word onset, and average power changes per subject were computed separately for referentially coherent and ambiguous trials.

### Statistical analysis

For each experiment, statistical evaluation of the time-frequency responses was performed with a cluster-based random permutation test (Maris and Oostenveld 2007). This statistical analysis was performed separately for the lower and higher frequency bands (2-30 Hz and 30-90 Hz, respectively).

For Experiment 1, analysis was performed in the 200 to 1500 ms latency window after onset of the pronoun (thereby excluding data associated with sentence-final words). For every data point (electrode by time by frequency) of the two conditions, a simple dependent-samples *t* test was performed (giving uncorrected *p*-values). All adjacent data points exceeding a pre-set significance level (5%) were grouped into clusters. For each cluster the sum of the *t* statistics was used in the cluster-level test statistic. Subsequently, a null distribution that assumes no difference between conditions was created. This distribution was obtained by randomly assigning the conditions in the subjects for a total of 1000 times and calculating the largest cluster-level statistic for each randomization. Finally, the actually observed cluster-level test statistics were compared against the null distribution, and clusters falling in the highest or lowest 2.5th percentile were considered significant (using the correcttail-option to correct *p*-values for doing a two-sided test).

For Experiment 2, the analysis was identical to the analysis for Experiment 1 but performed in a shorter 200 to 1000 ms latency window after onset of the noun phrase to avoid data associated with the disambiguation (average distance between onset of the noun phrase and onset of the disambiguating word was 1083 ms).

For Experiment 3, the analysis was identical to the analysis for Experiment 1 and 2 but performed in a 200 to 1000 ms latency window after onset of the determiner (which was also the window of analysis in Martin et al. 2012).

In Experiment 4, analysis was performed in a 200 to 1000 ms latency window after onset of the pronoun and was therefore identical to the analysis for Experiment 2 and 3.

### Beamformer source localisation of gamma-band activity (Experiment 4)

The data from Experiment 4 was collected from 64 EEG channels and because the larger number of electrodes allows a more reliable estimation of underlying sources, this experiment was best suited to attempt source localisation of the observed effects. We performed this source reconstruction with a beamforming approach, namely Dynamic Imaging of Coherent Sources (DICS; Gross et al. 2001), which uses an adaptive spatial filtering procedure to localize power in the entire brain.

Because referential coherence and ambiguity elicited significant differences in gamma activity around 40 Hz and around 80 Hz in different time intervals, we performed two separate source reconstructions. The first analysis contained data from 400-600 ms after CW onset in both conditions, therefore focusing on activity predominantly elicited by the pronoun itself. The second analysis contained data from 500-1000 ms after CW onset in both conditions. Both analyses involved comparison to the 500-300 ms pre-CW baseline period.

After extracting the data segments, all data was re-referenced to the average of all electrodes (common average reference), as is required for source localisation purposes. The first analysis centered on 40 Hz with frequency smoothing of ±5 Hz, and used a Hanning taper. The second analysis centered on 70 Hz with frequency smoothing of ±10 Hz, and used discrete prolate spheroidal sequences (Slepian sequences) as tapers.

Electrodes were aligned to a volume conduction model that was made based on a template brain using the boundary element method (Oostenveld, Praamstra, Stegeman, and van Oosterom 2001). A common spatial filter was then computed at 40 Hz or at 70 Hz for the pre-CW period and the post-CW period of both conditions together. The spatial filter was subsequently projected to all trials. Power values were calculated on an equidistant, three-dimensional template grid with a 5-mm resolution. Trials were then averaged in the pre-CW period and in the post-CW period separately, for each condition. Per condition, the power increase in the post-CW period relative to the pre-CW was computed in the following way: (post-CW – pre-CW) / pre-CW. After this, the grand averages were computed across subjects, and the difference between referential coherence and ambiguity was interpolated on the template brain for visualisation.

For the statistical analysis of the source reconstruction, we computed one-sided dependent sample t-statistics to compare the power values of the trial-averaged participant data of referential coherence and ambiguity at each of the 15,711 source points within the three-dimensional grid of the template brain. Clusters were constructed from source-locations with significant t-values of which neighboring locations also showed significant t-values. Then, the sum of the t-values in each cluster was calculated, and we selected the cluster with the largest sum of the t-values. For localizing the spatial coordinates of the significant areas, the t-values of the significant, clustered source points and zeros at all other points were interpolated to the template brain. We identified brain areas using a template atlas (Tzourio-Mazoyer et al. 2002).

## Results

In Experiment 1, we observed larger gamma-band power in the referentially coherent condition compared to the referentially ambiguous condition (*p* < 0.01), most prominent between 70 and 85 Hz and distributed from left-frontal to left-centroparietal electrodes (see Figure 1). No differences between the conditions were observed in the 2-30 Hz frequency band.

**Figure 1.**
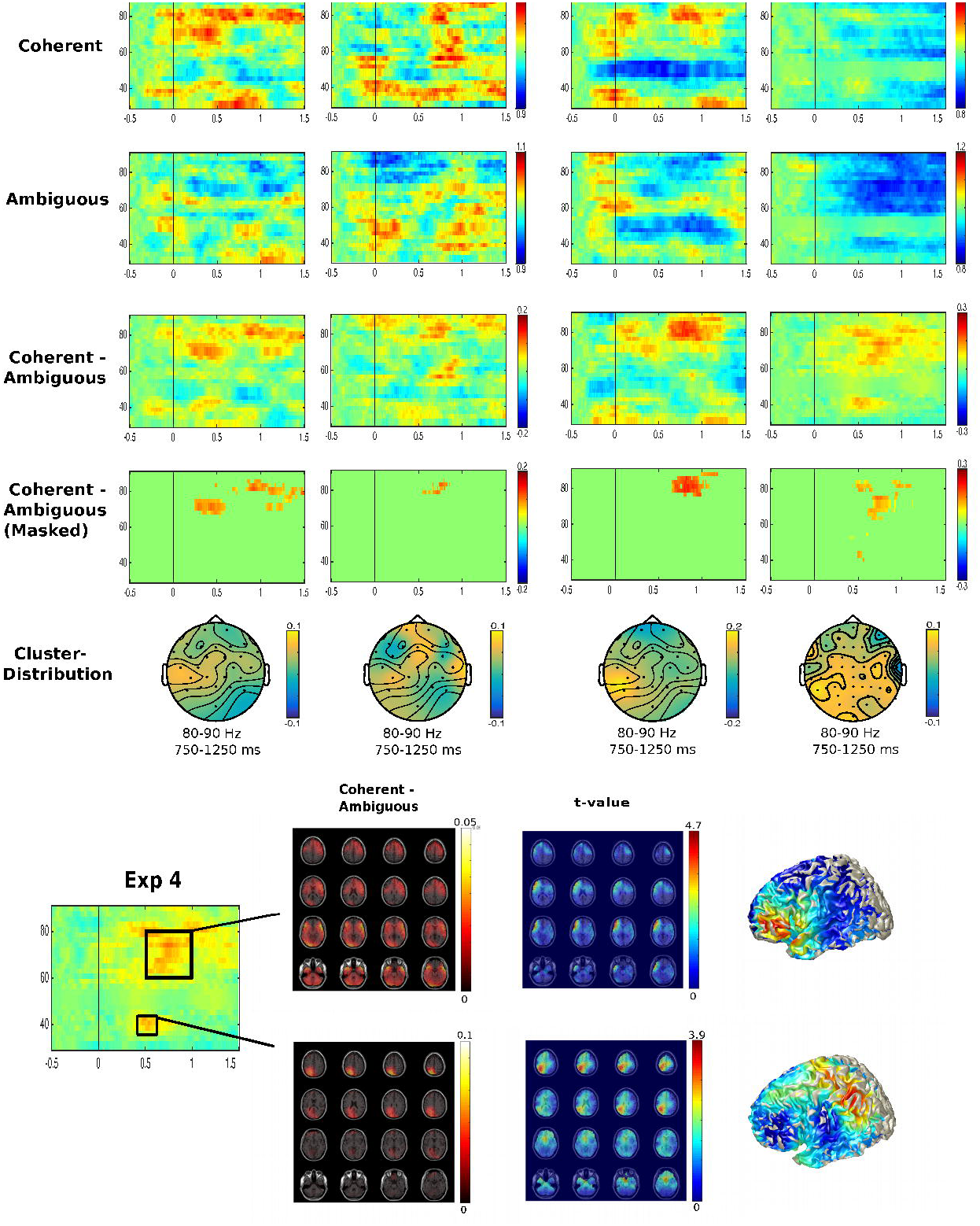
Grand-average oscillatory activity associated with referential coherence and referential ambiguity in each of the four experiments per column. The first and second row show oscillatory activity for the referentially coherent and ambiguous condition, respectively. The third row shows the condition difference (coherence minus ambiguity), and the fourth row shows that condition difference masked by the cluster permutation statistical results. The fifth row shows the scalp maps of the condition difference averaged in a frequency-band (80-90 Hz) and time window (750-1250 ms) that was representative for all four experiments. Source localisation results for Experiment 4 are shown in the bottom graphs (from bottom left to right: visualisation of the time/frequency data for the source localisation, slice plot view of the relative increase in gamma-band activity for referential coherence, slice plot view of the associated statistical results, surface plot view of the statistical results).

In Experiment 2, we also observed larger gamma-band power in the referentially coherent condition compared to the referentially ambiguous condition (*p* = 0.05), most prominent between around 80 Hz and distributed from frontal to centroparietal electrodes. As in Experiment 1, no differences between the conditions were observed in the 2-30 Hz frequency band.

In Experiment 3, we observed larger gamma-band power in the referentially coherent condition compared to the referentially ambiguous condition, most prominent between 75 and 85 Hz and distributed from left-frontal to left-centroparietal electrodes. Although this difference was only marginally significant (*p* < 0.1) using an unrestricted two-tailed cluster-permutation test, the difference was fully significant when taking into account the directionality of the effect expected based on Experiment 1 and 2 (i.e., a one-tailed test) or when restricting the analysis to a narrower gamma band (60-80 Hz) based on the results of Experiment 1 and 2. As in Experiment 1 and 2, no differences between the conditions were observed in the 2-30 Hz frequency band.

In Experiment 4, we observed larger gamma-band power in the referentially coherent condition compared to the ambiguous condition, most prominent between 60-80 Hz between approximately 500-1000 ms (*p* < 0.05), but also around 40 Hz in a second cluster between approximately 400-600 ms (*p* < 0.05). The bottom graphs in Figure 1 shows the visualisation plots of the source localisation results in the 40 Hz range (bottom-left graphs) and in the 70-90 Hz range (bottom-right graphs). The gamma-increase for referential coherence compared to ambiguity around 40 Hz was localised to left posterior parietal cortex (encompassing the superior parietal lobe and precuneus). The gamma-increase around 70 Hz was generated by left and right frontal-temporal regions (encompassing inferior frontal lobe, inferior temporal lobe and anterior temporal lobe), although the difference was only statistically robust in the left hemisphere.

Based on the observed gamma-band effect around 40 Hz in Experiment 4, we performed an additional frequency/time-of-interest analysis on the first three experiments. This analysis was a cluster-permutation test, as described earlier, restricted to average power values between 35-45 Hz in the 400-600 ms time window. Referential coherence was associated with significantly higher gamma-power than referential ambiguity in Experiment 1 (*p* < .05) and Experiment 2 (*p* = .05), but not in Experiment 3. Thus, although the unrestricted cluster permutation tests only revealed differences between coherence and ambiguity around 40 Hz in the Experiment 4, the more restricted follow-up tests revealed similar differences in Experiment 1 and 2.

## DISCUSSION

We conducted oscillatory analyses of four EEG experiments on reference resolution during language comprehension. We investigated the increased coupling of functional neural systems associated with referentially coherent expressions compared to referentially ambiguous or insufficient expressions, and the neural sources supporting these changes. Despite varying in modality, language and type of referential expression, all experiments showed larger gamma-band power for coherence compared to ambiguity. In Experiment 1, 2 and 4, a gamma-band increase was observed around 40 Hz around 400-600 ms after anaphor-onset and also at a higher frequency range (60-80 Hz) around 500-1000 ms. Beamformer analysis in high-density EEG Experiment 4 localised the increase around 40 Hz to left posterior parietal cortex and the increase in the 60-80 Hz range to left inferior frontal-temporal cortex. We argue that the observed gamma-band power increases reflect successful reference resolution, encompassing the reinstatement of antecedents by the brain’s recognition memory network and integration of antecedent information into the sentence representation by the frontal-temporal language network.

In the sections below, we first discuss our results in comparison to those of previous oscillatory studies on sentence- and discourse-level language comprehension, and we subsequently integrate our findings and previous results from patient and neuroimaging studies into a cortico-hippocampal theory of the neurobiology of reference. The corticohippocampal theory of reference is a first attempt to bridge three seemingly disparate areas of research, namely the cognitive/psycholinguistic study of reference, the neurobiology of recognition memory and the neurobiology of language. We discuss the predictions that follow from this theory and outline its major challenges for future research.

### Increased gamma-band activity for referential coherence

Our study is the first to investigate the neural oscillatory signature of successful reference, and generated a relatively robust pattern of results across four different experiments. Referentially coherent expressions led to increased gamma-band activity compared to referentially ambiguous expressions, and in three out of the four studies this activity occurred both at 40 Hz around 500 ms after onset and at 60-80 Hz around 500-1000 ms after onset. In conjunction with the source localisation results, we interpret these patterns to reflect the successful reactivation of antecedent information and integration into the unfolding sentence representation. For referentially ambiguous expression, no antecedent is reactivated to a sufficient degree and therefore, at least initially, no unique antecedent is successfully integrated into the unfolding representation. We thus assign two different functional interpretations for the 40 Hz and 70-80 Hz activity, although we acknowledge that the processes as reflected in these activity bands must work in close cooperation.

We note that our study is not the first to find gamma-band activity in these frequency-ranges modulated by a sentence-level linguistic manipulation. Several studies have reported increased gamma-activity for semantically coherent and predictable sentences compared to semantically anomalous or unexpected sentences (e.g., Wang et al. 2012; Rommers et al. 2013), and current debate centres on whether such patterns reflect effects of semantic unification/integration or effects of pre-activation/prediction (Wang et al. 2012; Lewis and Bastiaansen 2015a,b). We cannot logically rule out an explanation of our own findings in terms of prediction, as the observed effects may reflect the processing consequences of a successful prediction of a referential expression (see Bögels, Barr, Garrod and Kessler 2015, for a potential demonstration to this effect in a mentalizing task). However, we think that this explanation does not sit well with the absence of N400 ERP modulation in the four experiments that are analyzed here. In addition, our source localization results do not include the temporal lobe regions that show rapid effects of semantic predictability (e.g., Lau, Webder, Gramfort, Hämäläinen and Kuperberg 2013; Lau and Nguyen 2015), which typically show less activity to predictable words than to unpredictable words. For these reasons, an account of our results in terms of prediction is not particularly compelling.

Instead, we think that our 80 Hz gamma-findings are compatible with an interpretation in terms of ongoing sentence-level semantic unification operations, taking place predominantly in left inferior frontal cortex (e.g., Hagoort and Indefrey 2014). Such unification operations may proceed more fully and smoothly with a coherent anaphor than with an ambiguous or insufficient one. We also note the similarity of our 80 Hz findings in the left inferior frontal cortex and left anterior temporal lobe to results obtained with fMRI (Nieuwland et al. 2007, Figure 1E). In that previous study, coherent pronouns elicited stronger BOLD responses than ambiguous pronouns, particularly in left and right inferior frontal brain regions but with some extension into the anterior temporal lobe. We thus take these previous results and the current results as convergent evidence for the role of left inferior frontal regions in successful sentence-level unification operations.

### The cortico-hippocampal theory of reference

Our results are compatible with a novel neurobiological account that we dub the cortico-hippocampal theory of reference. This coarse-grained theory combines the cognitive architecture and processing principles from cue-based retrieval and psycholinguistic theories of anaphora (e.g., Dell, McKoon and Ratcliff 1983; Gernsbacher 1989; Sanford and Garrod 1989; Gerrig and McKoon 1998; McElree 2006; Martin 2016), with extant neurobiological theories of recognition memory (e.g., Rugg and Yonelinas 2003; Kahn, Davachi and Wagner 2004; Aggleton and Brown 2006; Eichenbaum, Yonelinas and Ranganath 2007) and of language comprehension (e.g., Price 2012; Friederici 2012; Bornkessel-Schlesewky and Schlesewsky 2013; Hagoort 2013; Hagoort and Indefrey 2014; Fedorenko and Thompson-Schill 2015; Friederici and Singer 2015;). Its core claim is that anaphor comprehension draws on the interaction between the recognition memory network (medial temporal lobe, including the hippocampus, and posterior parietal cortex) and the canonical frontal-temporal network, with the former being primarily responsible for reinstatement of the antecedent information and the latter being primarily responsible for integrating antecedent information into the unfolding sentence representation.

The currently available neurobiological and neuropsychological evidence in support of this theory is (1) observed BOLD increases in left-hippocampus for pronouns that uniquely match an antecedent compared to pronouns that match no antecedents (Nieuwland et al. 2007), (2) impairments in reference processing in patients with episodic memory dysfunction (hippocampal amnesia, Kurzcek et al. 2013; Alzheimer’s disease, Almor et al. 1999), and (3) the current findings of increased oscillatory activity in posterior parietal cortex and inferior frontal/temporal cortex for coherent reference.

The cortico-hippocampal theory of reference harbours at least the following predictions for future research. First, referential manipulations that compare old/new information (for example, anaphoric expressions compared to expressions that introduce a new discourse referent) elicit stronger activity difference in the recognition network than manipulations of the ease with which an anaphor is understood (e.g., a distance manipulation). Second, activity in the recognition network is of shorter duration, and may even take place before activity in the language network, reflecting the processing phases of antecedent reactivation and sentence-level integration (e.g., Sanford and Garrod 1989; Garrod and Terras 2000). Third, coherent reference should evoke stronger connectivity within the recognition network (e.g., between hippocampus and posterior parietal cortex) and between the recognition network and the language network. This connectivity could manifest itself in connectivity measures of BOLD fMRI, and possibly in the coupling of oscillatory activity (cross-frequency coupling). Fourth, reference to things that are perceptually available (e.g., visually present objects) do not engage the recognition network, but, instead may require multimodal integration through cooperation between the language network and brain regions involved in perceptual processing (see Martin 2016 for a sensory integration-based process model).

Beyond these predictions, there are also several important challenges to the corticohippocampal theory of reference. One is to determine the respective contributions of the medial temporal lobe and of posterior parietal cortex, and also of the various structures within these regions (e.g., Staresina, Fell, Lam, Axmacher and Henson 2012; Staresina, Cooper and Henson 2013; Staresina, Fell, Dunn, Axmacher and Henson, 2013). This is, of course, just as much a challenge to neurobiological theories of recognition memory as it is to the corticohippocampal theory of reference. An interesting possibility is that the hippocampus is initially involved in binding operations (e.g., Staresina and Davachi 2009) that link the anaphor to antecedent information (for a possible role of hippocampal binding in online language processes, see Duff and Brown-Schmidt 2012). Another challenge, in the frequency domain, is whether there is evidence for involvement of the well-established hippocampal theta rhythm and perhaps its coupling with gamma-band activity (e.g., Canolty et al. 2006; Jensen and Colgin 2007; Colgin et al. 2009; Canolty and Knight 2010). In the current study, we did not observe any hippocampal-theta activity changes as a function of referential coherence. However, hippocampal theta may not be reliably detectable in 64-channel surface-EEG data (e.g., Lopes da Silva 2013).

## CONCLUSION

We report time-frequency analysis of four EEG experiments to reveal the increased coupling of functional neural systems associated with referentially coherent expressions compared to referentially ambiguous expressions. Despite varying in modality, language and type of referential expression, all experiments showed larger gamma-band power for coherence compared to ambiguity or insufficient reference. We localised this increase in Experiment 4 to posterior parietal cortex around 400-600 ms after anaphor-onset and to frontal-temporal cortex around 500-1000 ms. The current findings can be synthesized with previous results from patient and neuroimaging studies such that, together, the evidence suggests a cortico-hippocampal neurobiological theory of reference. This corticohippocampal theory posits that anaphor comprehension draws on the interaction between the recognition memory network (medial temporal lobe and posterior parietal cortex) and the canonical frontal-temporal language network, with the former being primarily responsible for reinstatement of the antecedent information and the latter being primarily responsible for integrating antecedent information into the unfolding sentence representation. This neurobiological implementation would in turn offer support for a process model that incorporates the basic representational and mechanistic principles from the cue-based retrieval framework for language processing (e.g., McElree et al. 2003; Lewis et al. 2006; McElree 2006; Martin 2016). Taken together, these are the important first steps towards a fully articulated neurobiological theory of language that captures reference.

## ACKNOWLEDGEMENTS

We gratefully thank Lin Wang for making her data analysis scripts available to us. AEM is supported by ESRC Future Research Leaders fellowship and grant ES/K009095/1.

## Footnotes

1 Because of the very slow nature of this ERP effect (<0.5Hz), and because we applied a strict (48 dB/octave) 1 Hz high-pass filter before the oscillatory analysis, obtained oscillatory results do not also reflect the spectral representation of the previously reported grand-average ERP waveforms.

